# Heterotrophic Thaumarchaeota with ultrasmall genomes are widespread in the ocean

**DOI:** 10.1101/2020.03.17.996280

**Authors:** Frank O. Aylward, Alyson E. Santoro

## Abstract

The Thaumarchaeota comprise a diverse archaeal phylum including numerous lineages that play key roles in global biogeochemical cycling, particularly in the ocean. To date, all genomically-characterized marine Thaumarchaeota are reported to be chemolithoautotrophic ammonia-oxidizers. In this study, we report a group of heterotrophic marine Thaumarchaeota (HMT) with ultrasmall genome sizes that is globally abundant in deep ocean waters, apparently lacking the ability to oxidize ammonia. We assemble five HMT genomes from metagenomic data derived from both the Atlantic and Pacific Oceans, including two that are >95% complete, and show that they form a deeply-branching lineage sister to the ammonia-oxidizing archaea (AOA). Metagenomic read mapping demonstrates the presence of this group in mesopelagic samples from all major ocean basins, with abundances reaching up to 6% that of AOA. Surprisingly, the predicted sizes of complete HMT genomes are only 837-908 Kbp, and our ancestral state reconstruction indicates this lineage has undergone substantial genome reduction compared to other related archaea. The genomic repertoire of HMT indicates a highly reduced metabolism for aerobic heterotrophy that, although lacking the carbon fixation pathway typical of AOA, includes a divergent form III-a RuBisCO that potentially functions in a nucleotide scavenging pathway. Despite the small genome size of this group, we identify 13 encoded pyrroloquinoline quinone (PQQ)-dependent dehydrogenases that are predicted to shuttle reducing equivalents to the electron transport chain, suggesting these enzymes play an important role in the physiology of this group. Our results suggest that heterotrophic Thaumarchaeota are widespread in the ocean and potentially play key roles in global chemical transformations.

**Importance:** It has been known for many years that marine Thaumarchaeota are abundant constituents of dark ocean microbial communities, where their ability to couple ammonia oxidation and carbon fixation plays a critical role in nutrient dynamics. In this study we describe an abundant group of heterotrophic marine Thaumarchaeota (HMT) in the ocean with physiology distinct from their ammonia-oxidizing relatives. HMT lack the ability to oxidize ammonia and fix carbon via the 3-hydroxypropionate/4-hydroxybutyrate pathway, but instead encode a form III-a RuBisCO and diverse PQQ-dependent dehydrogenases that are likely used to generate energy in the dark ocean. Our work expands the scope of known diversity of Thaumarchaeota in the ocean and provides important insight into a widespread marine lineage.

## Introduction

Archaea represent a major fraction of the microbial biomass on Earth and play key roles in global biogeochemical cycles (1, 2). Historically the Crenarchaeota and Euryarchaeota were among the most well-studied archaeal phyla owing to the preponderance of cultivated representatives in these groups, but recent advances have led to the discovery of numerous additional phyla in this domain (3, 4). Among the first of these newly described phyla was the Thaumarchaeota (5), forming part of a superphylum that, together with the Aigarchaeota, Crenarchaeaota and Korarchaeota, is known as TACK (6). Coupled with recent discoveries of other lineages such as the Bathyarchaeota and Asgard (7, 8), the archaeal tree has grown substantially. Moreover, intriguing findings within the DPANN group, an assortment of putatively early branching lineages, has shown that archaea with ultrasmall genomes and highly reduced metabolism are common in diverse environments (9, 10). Placement of both the DPANN and the TACK phyla within the Archaea remains controversial (11, 12) highlighting the need for additional genomes from diverse archaea.

The Thaumarchaeota are particularly important contributors to global biogeochemical cycles, in part because this group comprises all known ammonia-oxidizing archaea (AOA) (13)), chemolithoautotrophs carrying out the first step of nitrification, a central process in the nitrogen cycle. Deep waters beyond the reach of sunlight comprise the vast majority of the volume of the ocean (14); in these habitats Thaumarchaea can comprise up to 30% of all cells and are a critical driver of primary production and nitrogen cycling (15, 16). Although much research on this phylum has focused on AOA, recent work has begun to show that many basal-branching groups or close relatives of the Thaumarchaeota are broadly distributed in the biosphere and have metabolism distinct from their ammonia-oxidizing relatives. This includes the Aigarchaeota, as well as several early-branching Thaumarchaeota, which have been discovered in hot springs and deep subsurface environments (17–19).

In this study we characterize a group of heterotrophic marine Thaumarchaeota (HMT) with ultrasmall genome size that is broadly distributed in the deep ocean waters across the globe. We show that although this group is a sister lineage to the AOA, it does not contain the molecular machinery for ammonia oxidation or the 3-hydroxypropionate/4-hydroxybutyrate (3HP/4HB) cycle for carbon fixation, but instead encodes numerous pyrroloquinoline quinone (PQQ)-dependent dehydrogenases and a divergent form III-a RuBisCO. The reduced genomic and metabolic features of HMT genomes are similar in some ways to some DPANN archaea, suggesting that divergent archaeal lineages may have converged on similar traits. Our work describes a non-AOA Thaumarchaeota that is ubiquitous in the dark ocean and potentially an important contributor to nutrient transformations in this globally-important habitat.

## Results and Discussion

### Phylogenomics and Biogeography of the HMT

We generated metagenome assembled genomes (MAGs) from 4 hadopelagic metagenomes from the Pacific (bioGEOTRACES samples) and 9 metagenomes from 750-5000 m depths in the Atlantic (DeepDOM cruise). After screening and scaffolding the resulting MAGs, we retrieved 5 high-quality HMT Thaumarchaeota MAGs with completeness > 75% and contamination < 2% (Table 1, see Methods for details). All MAGs shared high average nucleotide identity (ANI), reflecting low genomic diversity within this group irrespective of their ocean basin of origin (99.6% ANI between the Pacific and Atlantic MAGs, minimum of 97% ANI overall). Using the Genome Taxonomy Toolkit (20, 21) we classified the MAGs into the order *Nitrososphaerales* within the class *Nitrososphaeria*, indicating their evolutionary relatedness to ammonia oxidizing archaea (AOA). We also performed a phylogenetic analysis of the HMT MAGs together with reference Thaumarchaeota and Aigarchaeota using a whole-genome phylogeny approach, which suggests placement of the HMT as a sister clade to the AOA (Fig. S1). The reference genome UBA_057, which was previously assembled from a mid Cayman rise metagenome as part of a large scale genomes-from-metagenomes workflow (22), also fell within the HMT group (Fig S1).

**Table 1.** Statistics for the HMT MAGs presented in this study.

| MAG ID | Size (kbp) | No.<br>Contigs | No.<br>Proteins | Completeness | Contamination | Estimated<br>Genome<br>Size (kbp) | % G+C<br>content |
| --- | --- | --- | --- | --- | --- | --- | --- |
| HMT_ATL | 837.8 | 18 | 951 | 98.1 | 1.46 | 841.9 | 32.5 |
| HMT_PAC | 812.1 | 98 | 968 | 96.1 | 0.97 | 836.7 | 32.7 |
| HMT_AAIW_ATL | 697.5 | 60 | 812 | 78.2 | 0.97 | 883.7 | 32.8 |
| HMT_NADW_ATL | 670.8 | 34 | 766 | 76.6 | 0 | 875.5 | 32.6 |
| HMT_AABW_ATL | 814.8 | 36 | 913 | 88.8 | 0.97 | 908.3 | 32.7 |

All HMT MAGs encoded a full length 16S rRNA gene, and we therefore constructed a phylogenetic tree based on this marker to examine if previous surveys had identified the HMT lineage (Fig S2). We found that the 16S rRNA sequences of the HMT MAGs are part of a broader clade that has been observed previously in the Puerto Rico Trench (23), Arctic Ocean (24), Monterey Bay (25, 26), the Suiyo seamount hydrothermal vent water (27, 28), the Juan de Fuca Ridge (29), and deep waters of the North Pacific Subtropical Gyre (26, 30), Ionian Sea (31), and North Atlantic (32). Previous work has referred to this lineage as the pSL12-related group (26) and noted that it forms a sister clade to the ammonia-oxidizing Thaumarchaeota, consistent with our 16S and concatenated marker protein phylogenies (Figs S1, S2). Three fosmids were previously sequenced from this lineage (28) (AD1000-325-A12, KM3-153-F8, and AD1000-23-H12 in Fig S2), and their corresponding 16S rRNA gene sequences have 85-95% identity to the 16S rRNA genes of our HMT MAGs. Moreover, we compared the amino acid sequences encoded in the fosmids and found they have 41-72% identity on average to those of our HMT MAGs, suggesting that considerable genomic variability exists within this group in different marine habitats. Overall, the occurrence of sequences from this clade in such diverse marine environments suggests that the HMT lineage represents a broadly-distributed marine group that, although observed in numerous previous studies, has remained poorly characterized.

To investigate the biogeography of HMT Thaumarchaeota we leveraged our genomic data to assess the relative abundance of this group in mesopelagic Tara metagenomes from 29 locations (depths 250-1000 m) and 13 metagenomes from the bioGEOTRACES samples and DeepDOM cruises (depths 750-5601 m) (Fig 1a). Reads mapping to the HMT Thaumarchaeota were detected in all Tara mesopelagic samples, demonstrating their global presence in waters of the Atlantic, Pacific, and Indian oceans (Fig 1a). The relative abundance of HMT MAGs were significantly higher in bioGEOTRACES or DeepDOM metagenomes sampled below 1000 m (Mann-Whitney U-test, p< 0.005), suggesting that this group may be more prevalent in bathypelagic or hadopelagic waters, but further work will be necessary to confirm this given the small number of metagenomes available from waters deeper than 1000 m. Given the dominance of chemolithoautotrophic AOA in the dark ocean, we sought to compare the relative abundance of HMT to AOA in metagenomic samples. To do this we assessed the number of metagenomic reads mapping to the HMT-specific RuBisCO large subunit protein using a translated LAST search and compared the results to the number of reads that mapped to a set of Thaumarchaeota AmoA proteins (see Methods for details), with abundances normalized by protein length. This approach estimated that HMT can reach abundances up to 6% those of their AOA relatives (Fig 2b). Given the dominance of AOA in the dark ocean, these results suggest that HMT are both globally distributed and numerically abundant in the dark ocean, particularly in depths >1000 m. A recent study identified a the presence of Thaumarchaeota closely related to the group we report here in Monterey bay (depths 5-500 m) and reported that it represented abundances < 0.5% of the total Thaumarchaeota population, further suggesting that this group comprises a relatively small fraction of total archaea in shallow waters.

**Figure 1.**
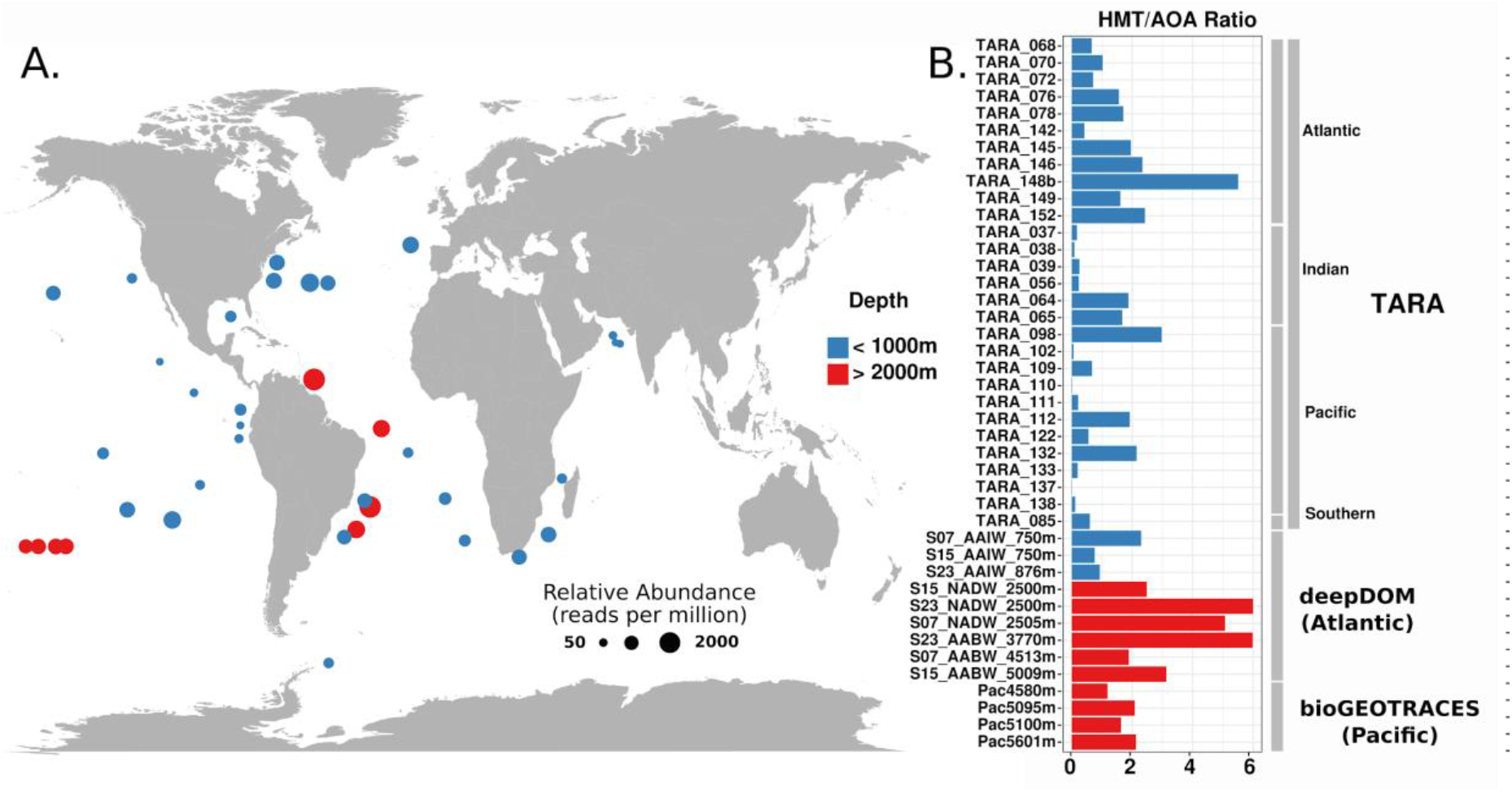
A) Relative abundance of the HMT lineage in metagenomic data collected from across the global ocean. Relative abundances were calculated by mapping metagenomic reads against a nonredundant set of HMT proteins, and are presented in units of reads per million. B) Relative abundances of HMT vs AOA were calculated by comparing the abundance of HMT RuBisCO to ammonia monooxygenase alpha subunit, and the values represent a percent of the AOA population (see Methods for details). Bars in red denote metagenomic samples taken at depths > 2000 m, while blue bars denote depths < 2000 m.

**Figure 2.**
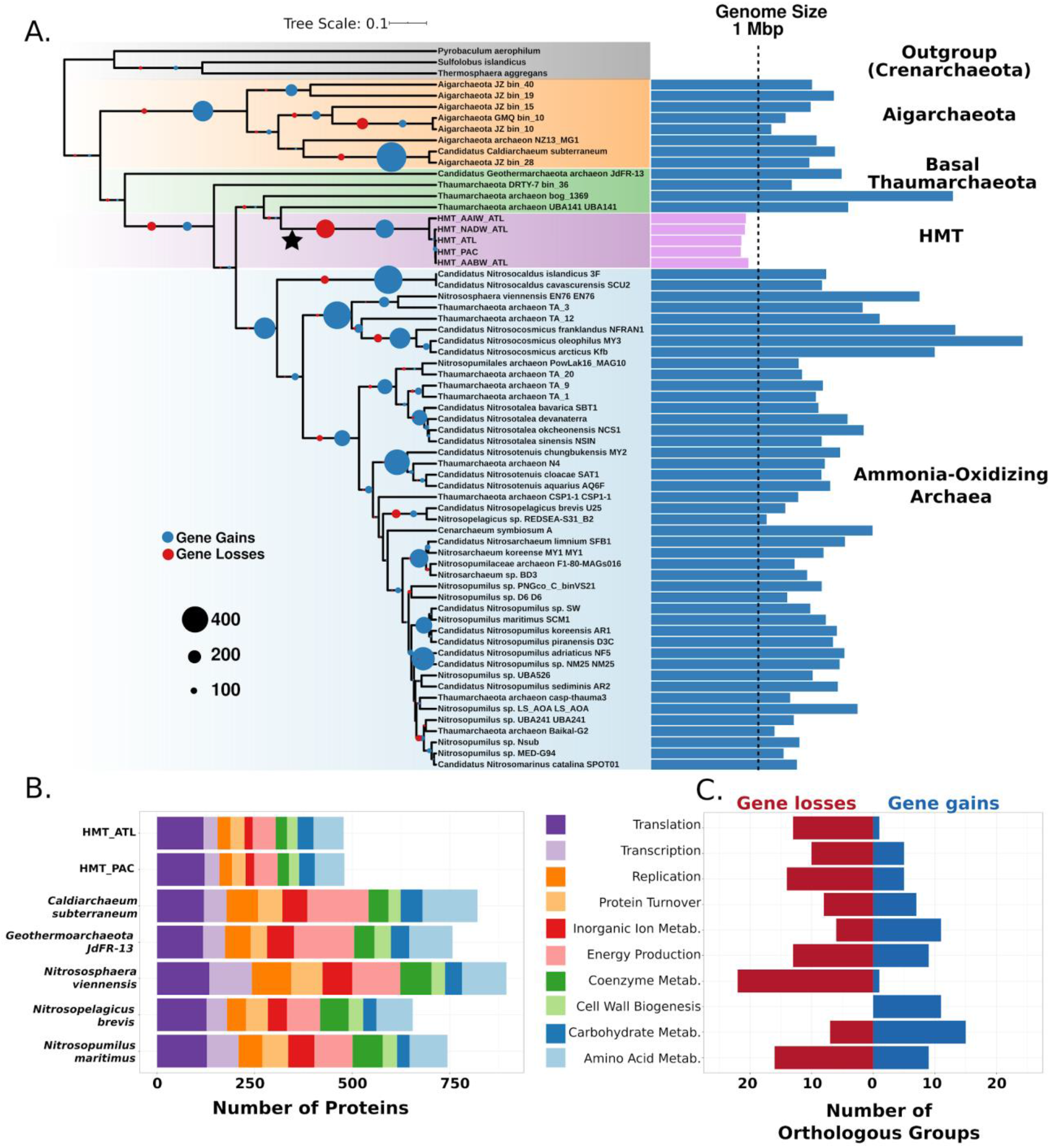
Phylogeny and Gene Loss Analysis. A) Phylogeny of high quality Thaumarchaeota genomes with genome sizes provided on the right. HMT genome sizes are colored purple. Complete genome sizes were estimated for MAGs using completeness and contamination estimates. Circles at the nodes provide the number of estimated gains and losses of orthologous groups. B) COG composition of two HMT genomes and select high quality reference Thaumarchaeota. Only genes annotated to select categories are provided; full annotations for all genomes are available in Dataset S3. C) Functional analysis of the OGs gained and lost on the branch leading to the HMT (star in panel A).

### Genome Reduction in the HMT lineage

The HMT MAGs were all between 671-838 Kbp in size, with 33% G+C content (Table 1). By extrapolating complete genome sizes using completeness and contamination estimates, we predict the complete genome sizes of this group are between 837-908 Kbp. This indicates that the HMT contain ultrasmall genomes that are smaller than any previously reported Thaumarchaeota, but similar to the size of some recently-described DPANN Archaea (9, 33). Small genome size often corresponds to small cell size in marine Bacteria and Archaea (34), and so the HMT Thaumarchaeota potentially have cell sizes smaller than the 0.15-0.2 μm diameter and 1-1.5 μm length range reported for marine AOA (35, 36). Given that some marine AOA can pass through the typical 0.2 μm pore size used for isolating cells from marine samples (36), it is possible that the HMT cells have eluded detection to date because they are not readily captured using this approach.

We calculated orthologous groups (OGs) between the HMT MAGs and a set of high-quality reference archaeal genomes (estimated completeness > 90% and contamination < 2%), resulting in total of 26,772 OGs (Dataset S1). We then performed ancestral state reconstructions on the OGs to distinguish between those that were lost by the HMT and those that were gained by other lineages. This analysis estimated that 197 OGs were lost on the branch leading to the HMT (Fig 2a), which is the largest single incidence of gene loss in our analysis. Compared to other reference Thaumarchaeota genomes, the HMT encoded markedly fewer genes involved in several broad functional categories, including energy metabolism, inorganic ion metabolism, coenzyme metabolism, and amino acid metabolism (Fig 2b). Moreover, many genes in these categories appear to have been lost specifically in the HMT lineage, indicating this group has undergone marked genome reduction, similar to many other marine lineages of Bacteria and Archaea (34). The HMT genomes lacked several highly conserved metabolic genes, including succinate dehydrogenase subunits, genes in tetrapyrrole biosynthesis (*hemABCD*), riboflavin metabolism (*ribABD, ribE, ribH*), and the genes involved cobalamin biosynthesis. The absence of cobalamin biosynthesis is coincident with the acquisition of the vitamin B12-independent methionine synthesis pathway (*metE*), and is in sharp contrast to AOA, which have been postulated to be major drivers of the production of this coenzyme (37, 38). Although these findings indicate that the HMT genomes have gone through a period of marked genome reduction, several genes have also been acquired in this lineage, most notably those involved in carbohydrate metabolism and cell wall biogenesis (Fig 2c), including several glycosyl transferases, pyrroloquinoline-quinone (PQQ)-dependent dehydrogenases, a phosphoglucosamine mutase, and several sugar dehydratases.

Unlike the ultrasmall genomes of the bacterial Candidate Phyla Radiation (CPR) and DPANN lineages (9), HMT genomes lack evidence to suggest they are merely DNA-containing intracellular components of larger cells. The HMT genomes contain an FtsZ-based cell division system, as well as components of Cdv-based cell division cycle. Genes for the synthesis of all amino acids are present in the same complement as other marine Thaumarchaeota (36), with exception of an alternate lysine biosynthesis pathway found primarily in methanogens (39). Though knowledge of the biosynthetic pathway for crenarchaeol and other glycerol dibiphytanyl glycerol tetraethers (GDGT) membrane lipids is incomplete, HMT appear to have the same identifiable components (e.g. geranylgeranyl reductase) present in other Thaumarchaeota. Together, this genomic evidence suggests that the HMT have retained a free-living lifestyle despite their pronounced genome reduction.

### Predicted Metabolism of the HMT lineage

HMT appear to be aerobic heterotrophs, in contrast to the AOA, as evidenced by the presence of the pyruvate dehydrogenase complex, a near complete TCA cycle, and full glycolysis pathway. No glycoside hydrolases were detected, suggesting they do not metabolize complex polysaccharides, but they do encode several ABC sugar transporters that may uptake simple oligo- and mono-saccharides (*malK, msmX, smsK, smoK, aglK, msiK*) or amino acids (*livFGHKM*), suggesting these as potential carbon sources. The HMT also encode a full Complex I (NUO operon) for the generation of a proton motive force, and cytochrome c as terminal oxidase. Like the AOA (40), HMT appear to utilize unusual blue-copper domain containing plastocyanins for electron transport with five such genes in the HMT_ATL MAG. There is no evidence that the HMT can use other terminal electron acceptors, as they lack nitrate, nitrite, and sulfate reductases. We also identified transporters predicted to target ammonia and phosphonates, suggesting sources of nitrogen and phosphorus, respectively. Interestingly, we did not identify any transporters predicted to target inorganic phosphorus, suggesting phosphonates may be the primary source of phosphorus for HMT.

The HMT genomes encode a single A-type ATPase similar to that encoded in neutrophilic AOA. A recent study showed that horizontal transfer has shaped the distribution of H+-pumping ATPase operons in Thaumarchaeota, with some deepwater or acidophilic Thaumarchaeota convergently acquiring a distinct V-type-like ATPase that potentially provides a fitness benefit in extreme environments (41). We performed a phylogenetic analysis of subunits A and B of this complex that demonstrated it has a phylogenetic signal consistent with the species tree for this group, with the HMT forming a sister clade to neutrophilic AOA (Fig S3). This indicates that despite the unusual genomic features of this group, the ATPase of the HMT has not been shaped by horizontal transfer in the same way as some hadopelagic or acidophilic Thaumarchaeota, and likely functions analogously to that of neutrophilic AOA.

Given the highly reduced genomes of the HMT lineage, it is notable that they encoded multiple PQQ-dependent dehydrogenases (quinoproteins), which are rarely found in archaeal genomes. Individual MAGs encoded between 4-9 individual PQQ-dependent dehydrogenases, and we cross-referenced all sequences to identify 13 total distinct copies, which we refer to here as PQQ dehydrogenase families. Additionally, two other HMT proteins are possibly members of this family based on their matches to the COG4993 glucose dehydrogenase family in the EggNOG database (AAIW_57_26 and AABW_353_2), but these proteins were not considered further since they did not match to any known Pfam families for these enzymes and they were considerably shorter than expected (less than 300 aa compared to > 600 aa for the other enzymes). PQQ-dependent dehydrogenases potentially target diverse carbon compounds and deliver reducing equivalents directly to the electron transport chain (ETC) through ubiquinone or cyctochrome c (42–44) without the need for energetic transport across the inner membrane (45), which is potentially of great importance in energy-limiting environments in the dark ocean. An analogous use of alcohol dehydrogenases to support ATP synthesis, but not biomass production, from diverse alcohols was shown for members of the SAR11 clade of alphaproteobacteria (46), also abundant in low nutrient environments. The HMT also encode a PQQ synthase, and can likely produce the cofactor for these enzymes, as well as a predicted cytochrome c oxidase subunit III, which potentially interacts with the PQQ alchohol dehydrogenases and facilitates their integration into the ETC. Moreover, our ancestral state reconstructions predict that this cytochrome c oxidase subunit was acquired in the HMT lineage along with the PQQ-dependent dehydrogenases (Fig. 2).

The divergent nature of the HMT quinoproteins makes substrate prediction for these enzymes extremely speculative. The most well-characterized of the PQQ-dependent enzymes are the membrane-bound glucose dehydrogenases (mGDH) and soluble methanol dehydrogenases (MDH). Both mGDH and alcohol dehydrogenases can act on a range of hexose and pentose sugars (47), with substrate flexibility potential determined by opening of the active site in the β-propeller fold (48). We note that all HMT quinoproteins lack the disulphide cysteine-cysteine motif common to all bona fide MDH (45, 49), but that 3 variants do retain a Asp-Tyr-Asp (DYD) motif common to the lanthanide-dependent methanol dehydrogenases and absent in other MDH forms (44), while another 4 retain a DXD motif without the conserved tyrosine (Fig. 3). Of the thirteen putative PQQ-dehydrogenase families, five contain signal peptides and twelve contain predicted transmembrane domains, suggesting membrane localization (Fig. 3). Moverover, Family 9 is represented by a single truncated enzyme encoded at the end of a contig (ATL_15_9), and it is therefore possible that a transmembrane domain is present in the full sequence. It is remarkable to note that 13 full-length PQQ-dependent dehydrogenases would require ~27 Kbp to encode, indicating that ~3% of the total genomic repertoire of the HMT is devoted to these enzymes alone. The large number of PQQ-dependent dehydrogenases together with the potential broad substrate specificity of these enzymes suggests that the HMT target a diverse range of compounds and that this is a key component of their metabolism.

**Figure 3.**
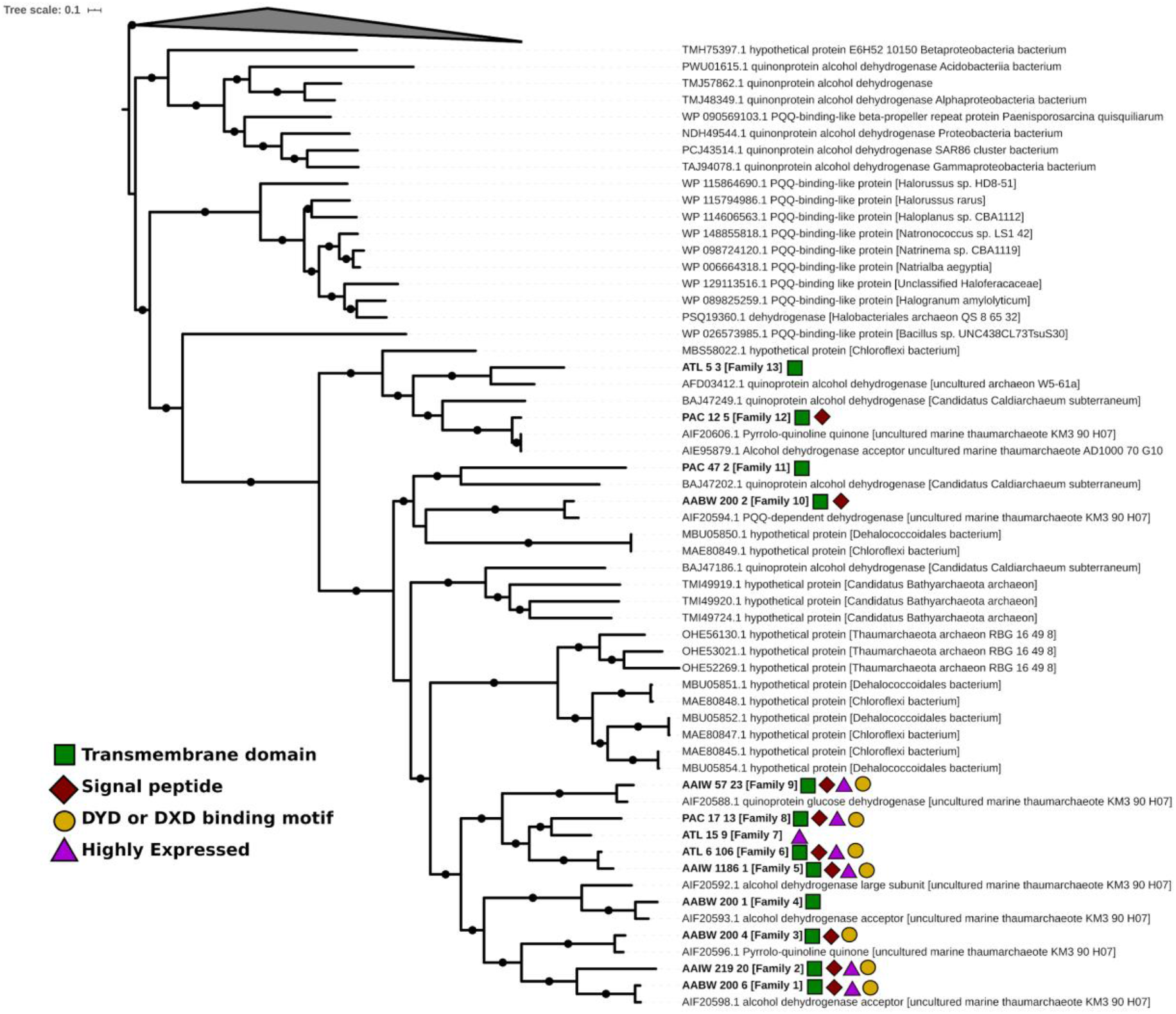
Maximum-likelihood phylogeny of 13 PQQ-dependent dehydrogenase families identified in the HMT genomes (bold) and reference sequences in NCBI GenBank identified with PSI-BLAST (101). Colors symbols indicate the presence of various features in the HMT enzymes. Enzymes that were among the top 30 most highly expressed genes in the DeepDOM metatranscriptomes are denoted with a purple triangle. A low confidence signal peptide was detected in the Family 13 PQQ-dependent dehydrogenase (ATL_5_3; 0.33 probability according to SignalP 5.0), which is not shown here. Black circles denote nodes with > 80% bootstrap support. The collapsed node contains 445 divergent PSI-BLAST hits that were used to root the tree.

The PQQ dehydrogenase families present in the HMT genomes are all divergent from references available in GenBank, and they share only between 31-91 % amino acid identity to one another. A phylogenetic analysis of these families together with available references, indicated that families 1-9 cluster together in a smaller sub-clade, while families 10-13 were more divergent (Fig. 3). Available reference proteins that clustered near families 10-13 were encoded within the Bathyarchaeota, Caldiarchaeum, and uncultivated Thaumarchaeota lineages, suggesting these enzymes may represent remnants of ancient enzymes that have long been present in the TACK superphylum. *Chloroflexi, Dehalococcoidales*, and *Bacillus* genomes also appear to encode related proteins, however, suggesting that horizontal gene transfer has also shaped the distribution of these enzymes to some extent. The family 1-9 enzymes clustered together with proteins encoded on fosmids sequenced from marine Thaumarchaeota in deep waters of the Ionian Sea (50) (KM3_90_H07 accessions in Fig. 3), suggesting that all of these enzymes are encoded within genomes of the same broad lineage. The clustering of the family 1-9 family enzymes together with fosmids from related Thaumarchaeota suggests that these enzymes may have evolved through a process of duplication and divergence that took place prior to the diversification of these lineages. Thus, the large number of PQQ-dependent dehydrogenases present in the HMT genomes may be due to lineage-specific expansion of this protein family in the distant past that occurred concomitant with adaptation to their current ecological niche.

The presence of a large subunit RuBisCO homolog (*rbcL*) in the HMT genomes is a notable feature of this group. We identified this gene in 4 of 5 HMT MAGs (missing only in HMT_AABW), but given the high nucleic acid similarity within the HMT it is likely that it is encoded in all genomes within this group but was merely not assembled in one MAG. We performed a phylogenetic analysis of the HMT RuBisCO homologs that placed it with other form III-a members of this protein family, albeit with long branches that demonstrate it is divergent from any previously-identified homolog (Fig S4a). To assess the likelihood that the HMT RuBisCO homologs are functional, we searched for 19 residues that have been shown to be critical for the substrate binding and activity of this enzyme (51, 52), and successfully identified conservation of 17 of these residues (Fig S4b). The two positions in the alignment where conserved residues were not conserved were 223 (aspartate instead of glycine), and 226 (tyrosine instead of phenylalanine or another hydrophobic residue), but a recent motif analysis of the RuBisCO protein family has shown that two positions tend to be among the most variable of the 19 conserved residues (53), indicating that activity may be retained despite these differences. Several recent studies have identified RuBisCO in disparate archaeal lineages; one study of Yellowstone hot spring metagenomes provided the first report of RuBisCO-encoding Thaumarchaeota (Beowolf and Dragon Archaea) (17), and the authors of this study postulated that this enzyme may function as part of an AMP-scavenging pathway in these archaea. Several subsequent studies identified diverse RuBisCO in a large number of DPANN Archaea (9, 53), and these studies further postulated that many of these enzymes participate in nucleotide scavenging, with one even demonstrating the activity of a form II/III version of this enzyme (54). It is interesting to note that many of the RuBisCO-encoding DPANN also have ultrasmall genomes and reduced metabolism (33), similar to HMT, suggesting that these features appear to have evolved independently in disparate archeal lineages.

To provide additional insight into the metabolic priorities of the HMT lineage we analyzed their *in situ* gene expression patterns in 10 metatranscriptomes collected at depths 750-5000 m in the South Atlantic during the DeepDOM cruise (Fig. 4, Dataset S2, details in Methods). The PQQ-dependent dehydrogenases were among the most highly expressed genes across all samples, and seven had expression levels orders of magnitude larger than most other genes (denoted with purple triangles in Fig 3), further highlighting the important role of these enzymes in the physiology of HMT. Several transporters predicted to target ammonium, phosphonate, and amino acids, a ribosomal protein, chaperones, and several hypothetical proteins were also among the most highly expressed (Fig. 4, Dataset S2). To evaluate the consistency of HMT transcriptional profiles, we also examined HMT expression in 12 metatranscriptomes associated with water near hydrothermal vents of the Mid-Cayman Rise in the Caribbean Sea (55). Although HMT transcript abundance was lower in these samples, several PQQ-dependent dehydrogenases were once again the most highly expressed genes (Fig S5). Overall the transcriptional profiles were markedly similar (Pearson’s r = 0.81 between mean transcriptional profiles in both datasets), and only 30 genes were differentially expressed (Wald test in DESeq2, corrected p-value < 0.005), suggesting that the transcriptional activities of HMT does not vary dramatically between disparate deepwater habitats. Together with the highly reduced genomes and metabolism of HMT, the lack of complex transcriptional regulation is consistent with the hypothesis that this group has a simple and consistent heterotrophic lifestyle and does not markedly alter its metabolic state based on environmental stimuli.

**Figure 4.**
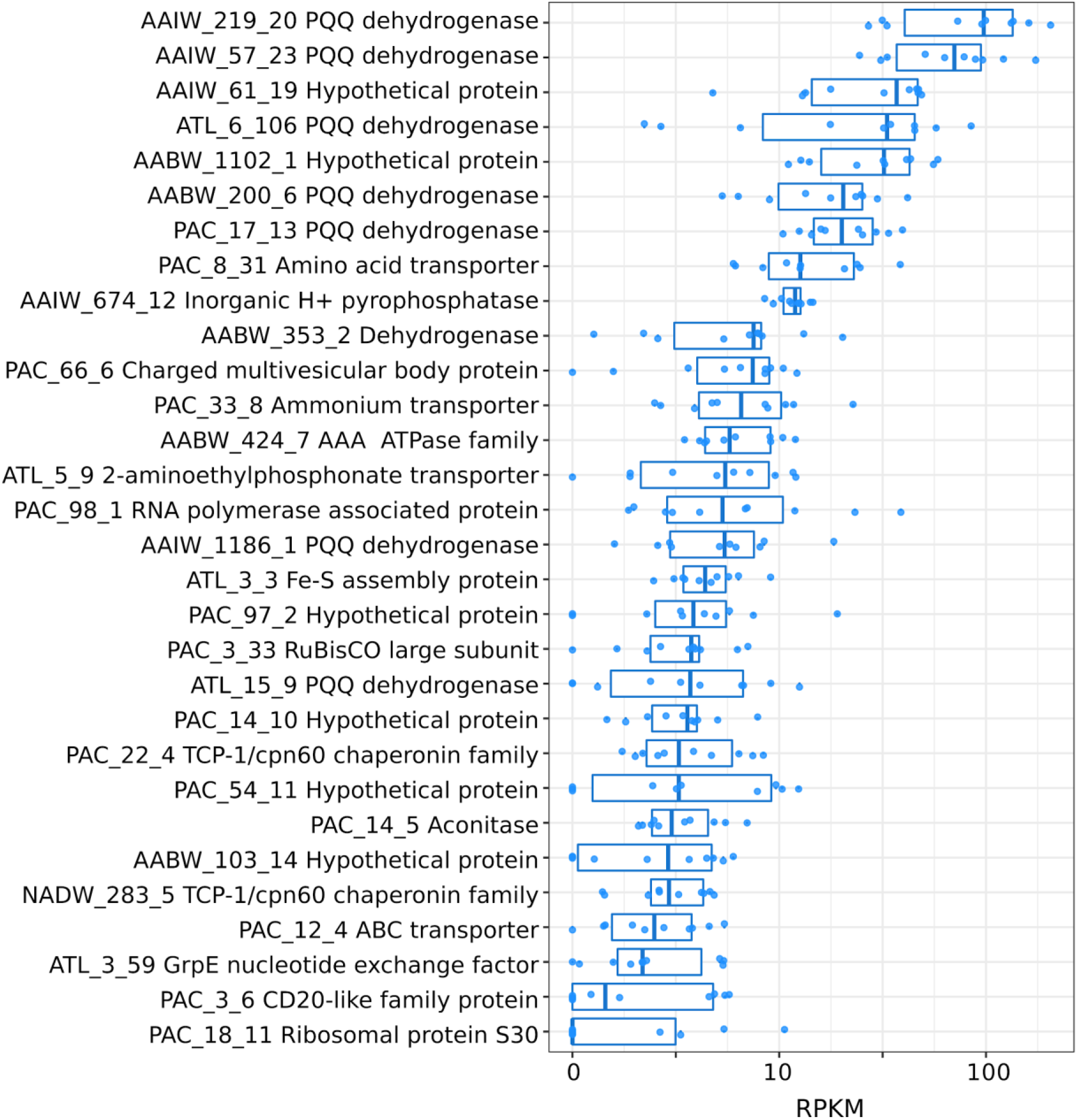
Top 30 HMT genes with highest median RPKM in 10 metatranscriptomes generated from South Atlantic waters at depths 750-5000m. Protein names refer to a non-redundant set of HMT proteins collectively present in the HMT MAGs.

### Conclusion

In this study we characterize a group of heterotrophic marine Thaumarchaeota (HMT) that is widespread in the global ocean. By analyzing MAGs from this group assembled from Atlantic and Pacific metagenomes we show that they have highly reduced genomes and a predicted heterotrophic metabolism. Paralogous expansion of PQQ-dependent dehydrogenases suggest the importance of oxidizing diverse carbon compounds and introducing reducing equivalents directly into the electron transport chain, which may be a critical component of their physiology in deep waters where energy is scarce. These PQQ dehydrogenases are among the most highly expressed genes in HMT and comprise up to ~3% of the total bp in their genomes, underscoring their likely importance. Further, HMT also encode a highly expressed form III-a RuBisCO that potentially functions as part of a modified nucleotide scavenging pathway, and therefore also benefits HMT in an energy limiting environment.

Ribosomal rRNA gene surveys have previously identified this group (26, 28), and closely related sequences are observed in diverse ocean provinces around the globe (23, 24, 27, 30–32). One study sequenced three fosmid sequences from the HMT group and noted that one even encoded a PQQ-dependent dehydrogenase (28), however, these fosmids only had 41-72 % amino acid identity and 85-95% 16S identity to our HMT genomes, and therefore likely represent a distinct group within this lineage. This suggests that the MAGs we present here, which all had 99% nucleic acid identity to each other, likely represent only a subset of the overall diversity of heterotrophic Thaumarchaeota in the ocean. Another recent study examining two MAGs assembled from Monterey Bay enrichment cultures reported a pSL12-like group of Thaumarchaeota with heterotrophic metabolism and a type III RuBisCO (56), and the 16S rRNA genes contained in these genomes cluster with our HMT MAGs (ASW8 sequence in Fig. S2). These similarities make it likely that the ASW8 MAG falls within the broader HMT lineage that we describe here, although it is unclear how much genomic heterogeneity exists within this group in nature, and further studies will be needed to evaluate the range of genomic diversity of HMT in marine environments.

Although previous work on marine Thaumarchaeota has been focused on chemolithoautotrophic ammonia-oxidizing lineages, our findings lead to the surprising conclusion that heterotrophic Thaumarchaeota are also widespread in the global ocean. In addition to broadening our understanding of archaeal diversity in the ocean, this finding could have multiple implications for biogeochemical cycling in the deep ocean. First, archaeal lipid distributions are used as paleoproxies of past ocean temperature. Current interpretations of the isotopic composition of archaeal lipids from marine sediments (e.g. (57)) and the water column (58) require up to 25% of archaeal carbon to be heterotrophic in origin, or invoke variable isotopic fractionation (59). If heterotrophic HMT are, as our data suggest, as much as 6% of the planktonic AOA community, this could provide some of the heretofore “missing” heterotrophic signal in the archaeal lipid data. Second, the quinoprotein-facilitated oxidation of organic carbon compounds consumes O_2_ if electrons are passed through the entirety of the ETC, but does not immediately yield CO_2_. If the products of these dehydrogenases are released and not further assimilated by the cell, this would lead to the consumption of deep ocean O_2_ that is not coupled to a corresponding consumption of dissolved organic carbon (DOC) and an anomalous respiratory quotient (60). Such cycling could provide a specific mechanism whereby DOC in the deep ocean decreases in lability with only minor changes in concentration (61). Further studies will be needed to examine the implications posed by the presence of these globally distributed heterotrophic Thaumarchaeota in the ocean.

## Methods

### Metagenomes

We constructed MAGs from metagenomic data generated from the DeepDOM cruise in the South Atlantic in April 2013 (62, 63) and bioGEOTRACES samples collected during the Australian GEOTRACES Southwestern Pacific section (GP13) in May-June 2011. Data from the bioGEOTRACES samples have been described previously (64), and only metagenomic datasets corresponding to depths > 4000 m were examined here. DeepDOM metagenomes were generated using methods similar to those previously described (65), and metagenomes were processed through the Integrated Microbial Genomes/Microbiomes (IMG/M) workflow at the Joint Genome Institute (66). Metagenome samples correspond to the IMG/M accessions 3300026074, 3300026079, 3300026080, 3300026084, 3300026087, 3300026091, 3300026108, 3300026119, and 3300026253. To generate a HMT MAG from the South Pacific, we co-assembled reads from four deepwater metagenomes from the bioGEOTRACES samples (64) (SRA accessions SRR5788153, SRR5788420, SRR5788329, SRR5788244).

### MAG construction

We constructed metagenome-assembled genomes (MAGs) using MetaBat v. 2.0 (67). We binned contigs in each metagenome using the following four different parameter sets: 1) -m 5000 -s 400000, 2) -m 5000 -s 400000 --minS 75, 3) -m 5000 -s 400000 --maxEdges 150 --minS 75, and 4) -m 5000 -s 400000 --maxEdges 100 --minS 75 --maxP 90. We evaluated the quality of the resulting MAGs using CheckM v 1.0.18 (68), and we then generated a set of de-replicated high quality MAGs using dRep v. 2.3.2 (69). Only MAGs that were estimated to be > 50% complete with < 2% contamination were considered further. Due to the short branch-length of many of the HMT genomes in our multi-locus tree, we postulated that most of these MAGs derived from the same population and could therefore be combined together to generate higher quality assemblies. To achieve this we scaffolded multiple MAGs together using Minimus2 in the AMOS package (70), ultimately producing the highest quality merged MAG by combining 2500_S23_NADW_med.34.fa and 5009_S15_AABW_high.53.fa (referred to as HMT_ATL, estimated to be 98% complete with 1.97% contamination). To generate the Pacific HMT MAG (HMT_PAC) we mapped reads from the four bioGEOTRACES samples mentioned above against the HMT_ATL MAG using bowtie2 v. 2.3.4.3 (default parameters, (71)), and subsequently pooled and re-assembled all mapped reads using SPAdes v. 3.13.1 (default parameters) (72). The resulting MAG, referred to as HMT_PAC, has estimated completeness of 96.13 % and contamination of 0.97% (Table 1).

### MAG Annotation

We predicted genes and proteins from the HMT genomes using Prodigal v 2.6.3 (73) and subsequently annotated the proteins by comparing them to the EggNOG 4.5 (74) and Pfam 31.0 (75) databases using the hmmsearch command in HMMER3 (76) (parameters -E 1e-5 and --cut_nc, respectively). Additionally, we retrieved KEGG annotations using the KEGG KAAS server (single direction method) (77). We predicted rRNAs on all MAGs using barrnap v. 0.9 (https://github.com/tseemann/barrnap). Signal peptides were predicted using the SignalP-5.0 server (78), transmembrane topology was predicted using Phobius (79), and transporters were analyzed using the TransAAP server in the TransportDB 2.0 utility (80). Estimated complete genome sizes were calculated from total bin sizes and contamination and completeness estimates using methods previously described (81). To generate a non-redundant set of HMT proteins for transcriptome mapping we used CD-HIT (default parameters) (82), resulting in a set of 1079 proteins. Annotations discussed in the text are largely based on the two best quality HMG MAGs (HMT_ATL and HMT_PAC), but full annotations were assessed for some enzymes such as the PQQ-dependent dehydrogenases. Annotations for all MAGs can be found in Dataset S3.

Some HMT genomes encoded PQQ-dependent dehydrogenases that were at the end of contigs and appeared truncated. To avoid mis-annotations based on protein truncation we compared these enzymes between all HMT MAGs and selected a set of best-quality representative proteins to use for detailed annotation and phylogenetic analysis (names provided in Fig. 3). The HMT_ATL MAG encoded one PQQ-dependent dehydrogenase that appeared truncated (ATL_15_9; 223 aa compared to > 600 aa for the others), but no similar homolog could be found in any of the other MAGs. We used ATL_11_9 for downstream annotations, but these should be treated with caution given this is likely not a complete protein sequence.

### Read mapping from TARA metagenomes

To estimate the global abundance and biogeography of HMT we mapped raw reads from mesopelagic samples from the Tara Oceans expedition (83) onto a nonredundant set of HMT proteins using LASTAL (84) (parameters -m 10, -Q 1, -F 15). Only hits having bit scores > 50 and % identity > 90 were retained. To compare the relative abundance of HMT and Ammonia oxidizing Archaea (AOA), we mapped reads to both the RbcL protein of the HMT_ATL and a selection of 31 AmoA genes from representative AOA in NCBI RefSeq, with only best hits retained. For both whole genome and marker gene read mapping we normalized the number of mapped reads by depth of sequencing to arrive at units of reads per million. For the RbcL/AmoA comparison we further normalized these relative abundances by the length of each reference protein to arrive at final units of reads per million per 100 amino acids, which we used for final comparisons. All values can be found in Dataset S4.

### Orthologous Groups

We predicted proteins from each of the HMT genomes using Prodigal v. 2.6.3 (73), and subsequently generated orthologous groups using Proteinortho v. 6.06 (85) (default parameters). We selected a representative protein from each OG at random and used these for subsequent annotations. We used the hmmsearch command in HMMER3 to compare these proteins to EggNOG 4.5 (e-value cutoff of 1e-5) and Pfam v. 31 (--cut_nc cutoffs), and the KEGG KAAS server to retrieve KO accessions, as described above for the genome annotations.

### Molecular Phylogenetics

We generated a multi-locus concatenated protein phylogeny of the HMT MAGs, representative Thaumarchaeota, and 3 Crenarchaeal outgroups using a set of 30 marker genes that includes 27 ribosomal proteins and 3 RNA polymerase subunits. We predicted marker proteins using the markerfinder.py script, which is available on GitHub (github.com/faylward/markerfinder). This script uses a set of previously described Hidden Markov Models to identify highly conserved marker genes in genomic data (86). We aligned marker proteins separately and trimmed the alignments using trimAl (87) (parameter “-gt 0.1) before inferring a maximum likelihood tree using IQ-TREE (88) with ultrafast bootstrap support used to infer confidence (89). The model LG+F+I+G4 was the best fit model inferred using the ModelFinder utility in IQ-TREE (90).

For the 16S tree, we obtained representative marine Thaumarchaeal sequences from NCBI GenBank (91) and the Ribosomal Database Project (RDP) (92). Reference sequences with high similarity to HMT 16S sequences were initially identified using the Classifier in RDP (93). We aligned reference and HMT 16S sequences using MAFFT (94) with the ‘--localpair’ option, trimmed the alignment with trimAl (-automated1 option), and generated the tree with IQ-TREE using the SYM+G4 model, broadly consistent with previous approaches (3).

### Transcriptome Mapping

We analyzed 10 metatranscriptomes collected during the same DeepDOM cruise in which the metagenomes were collected. These samples correspond to IMG/M accessions 3300011314, 3300011304, 3300011321, 3300011284, 3300011290, 3300011288, 3300011313, 3300011327, 3300011318, and 3300011316. Metatranscriptomes were processed consistent with methods previously described (65). In addition, to compare HMT expression patterns in a different deep sea environment we downloaded and mapped reads from 12 metatranscriptomes associated with hydrothermal vents (55). These samples correspond to NCBI Sequence Read Archive accessions SRR2044842, SRR2044843, SRR2044844, SRR2044846, SRR2044878, SRR2044888, SRR2044889, SRR2044891, SRR2044892, SRR2044911, SRR2044912, SRR2044914. We mapped reads from all metatranscriptomes onto a consolidated non-redundant set of HMT proteins (see MAG Annotation section above), using LAST with parameters “-m 10 -u 2 -Q 1 -F 15”, and the reference database was first masked with tantan (95), consistent with previously described methods (96). We only considered hits with bit scores > 50 and percent identity > 90%, and we processed LAST mapping outputs consistent with previous methods (97) and normalized transcript counts using the Reads per Kilobase per Million (RPKM) method (98). Detailed information can be found in Dataset S2. For differential expression analysis we used the DESeq2 package (99), with raw counts imported into a DESeq2 dataset and the 12 DeepDOM and 10 hydrothermal vent samples partitioned as conditions. Corrected p-values < 0.005 of the Wald test considered significant.

### Ancestral state reconstruction

We performed ancestral state reconstructions using the 5 HMT MAGs and a set of 60 high quality reference Thaumarchaeota genomes available in NCBI as of Dec. 1st, 2019. We only considered genomes with estimated completeness > 90% and contamination < 2%. The genomes were chosen to represent as broad a phylogenetic breadth of Thaumarchaeota as possible and without over-representing any particular phylogenetic group (for example, many high quality genomes of ammonia-oxidizing Thaumarchaea are available, but several of these were excluded if close relatives were still represented). We estimated the presence of OGs in internal branches of the Thaumarchaeota tree using the “ace” function in the “ape” package in R (100), with OG membership treated as a discrete feature. We constructed a binary matrix of extant OG membership from the Proteinortho output and subsequently inferred ancestral OG membership at each internal node on an ultrametric tree, which we constructed using the “chronos” function. We rounded log-likelihoods to the nearest integer to infer the probability of OG presence/absence at each internal node.

## Data Availability

Genomes for the HMT MAGs have been submitted to NCBI GenBank and accession numbers are pending. The genomes of the 5 HMT MAGs are also available on the Aylward Lab FigShare account: https://figshare.com/articles/HMT_genomes_tar_gz/11981253.

## Acknowledgements

We acknowledge the use of the Virginia Tech Advanced Research Computing Center for bioinformatic analyses performed in this study. This work was supported by grants from the Institute for Critical Technology and Applied Science and the NSF (IIBR-1918271), a Sloan Research Fellowship in Ocean Sciences, and Simons Early Career Awards in Marine Microbial Ecology and Evolution to FOA and AES. The DeepDOM cruise was supported by NSF OCE-1154320 to E. Kujawinski. Metagenomic and metatranscriptomic data from the DeepDOM cruise were generated under a US Department of Energy Joint Genome Institute Community Sequencing Program award to S. Hallam, E. Kujawinski, K. Longnecker and M. Bhatia. We thank them for permission to use the data in this manuscript.

## Figures

**Figure S1.**
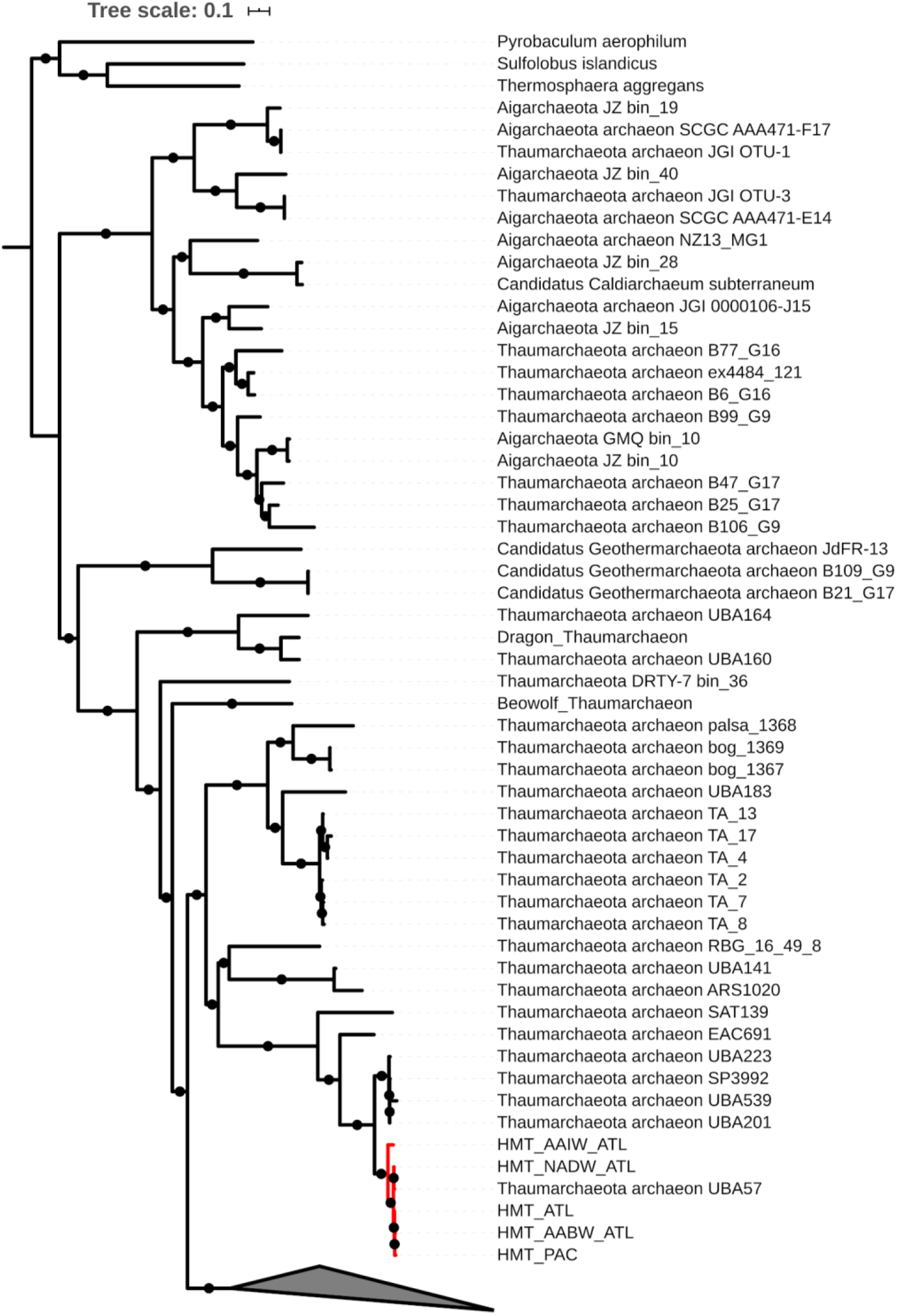
Concatenated phylogeny including Thaumarchaeota genomes with completeness > 50%. The HMT clade is shown in red. The collapsed node contains 147 genomes of Ammonia Oxidizing Archaea (AOA).

**Figure S2.**
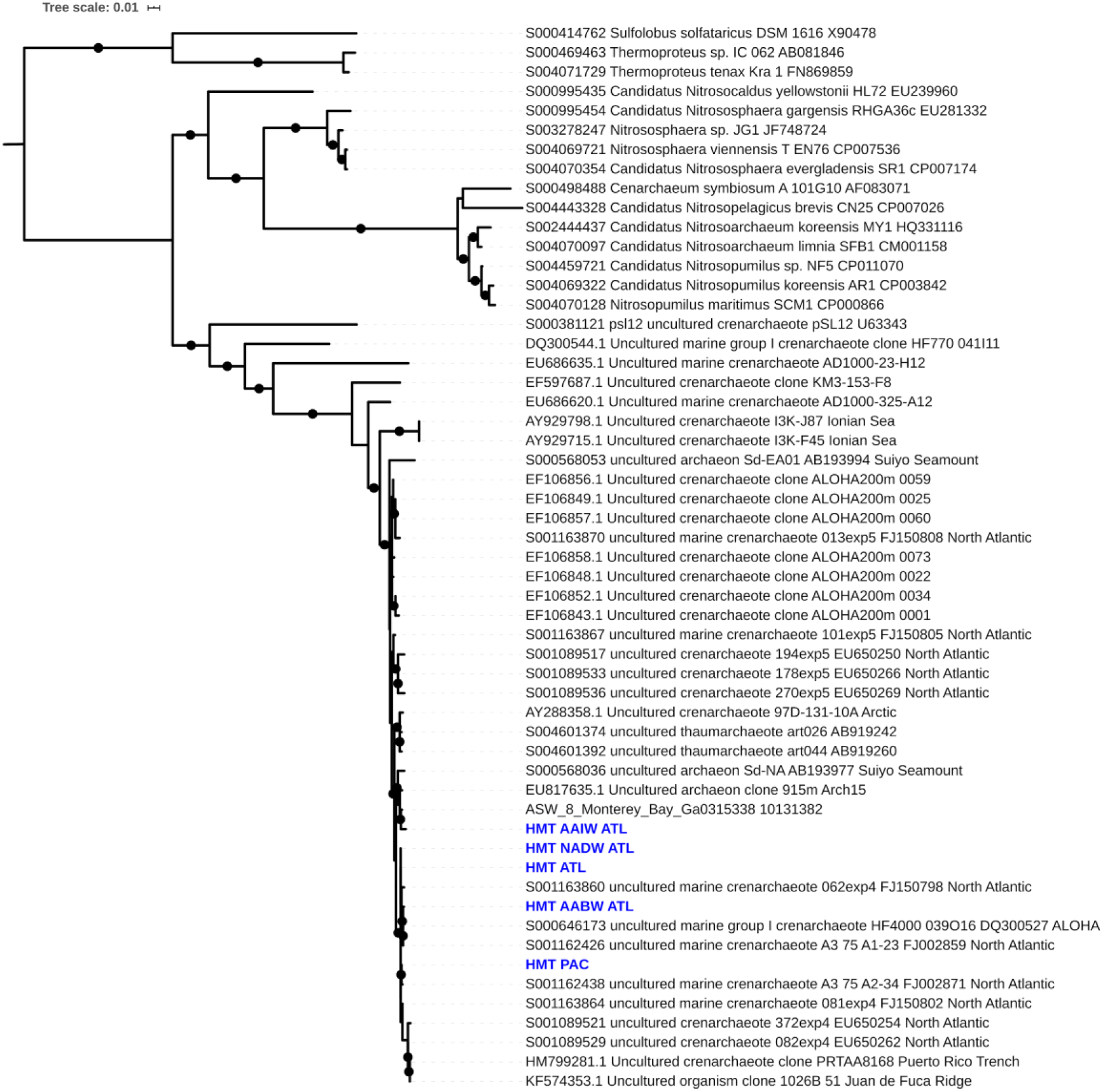
Maximum likelihood tree of the HMT using 16S rRNA genes and references available in NCBI and the Ribosomal Database Project. Nodes with ultrafast bootstrap support > 80 are denoted with a solid circle. 16S sequences from fosmids discussed in the text are bold, and HMT MAG 16S sequences are colored blue.

**Figure S3.**
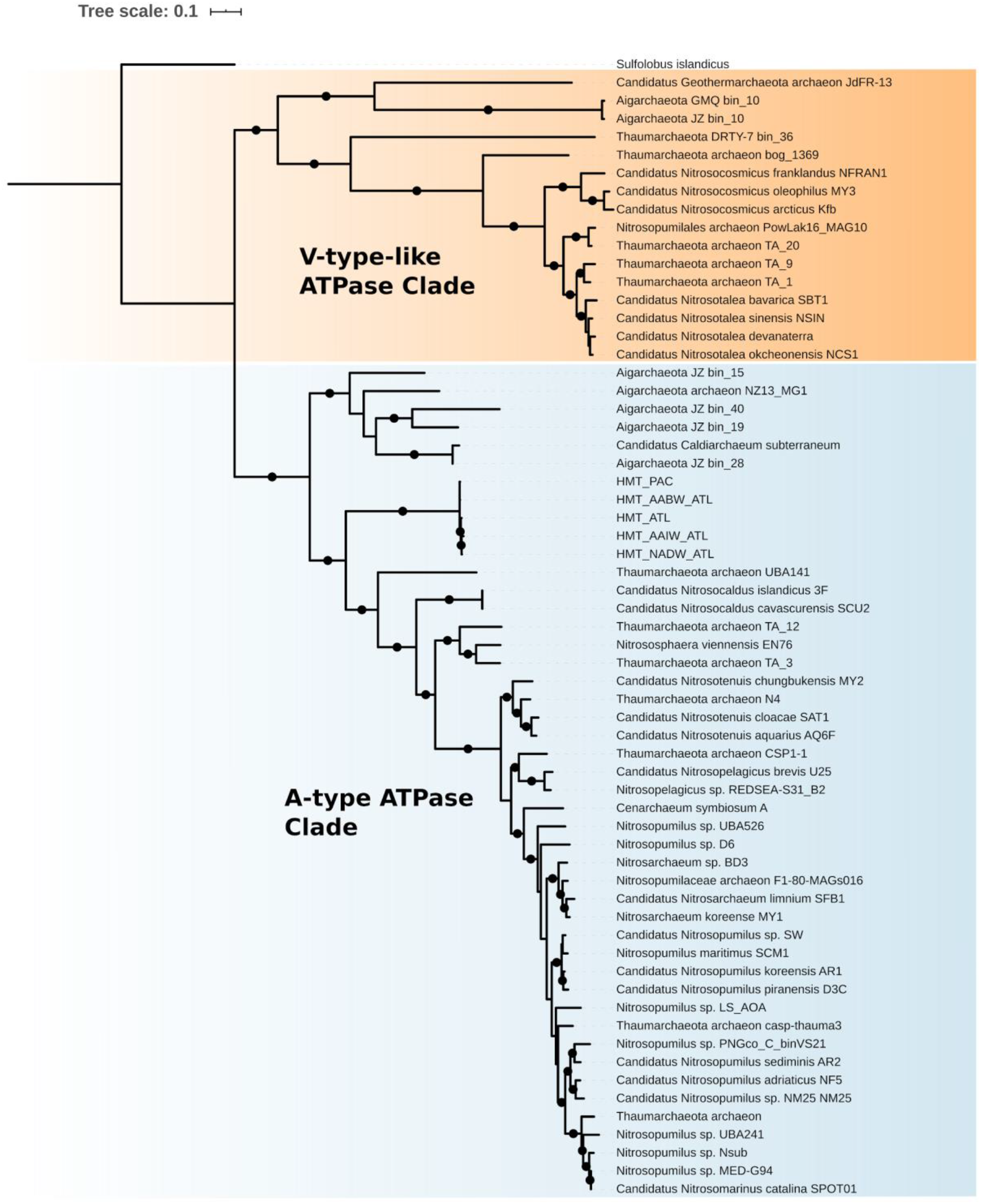
Maximum likelihood phylogeny of the ATPase subunits present in the HMT genomes and available Thaumarchaeota references. The phylogeny is based on a concatenated alignment of ATPase subunits A and B.

**Figure S4.**
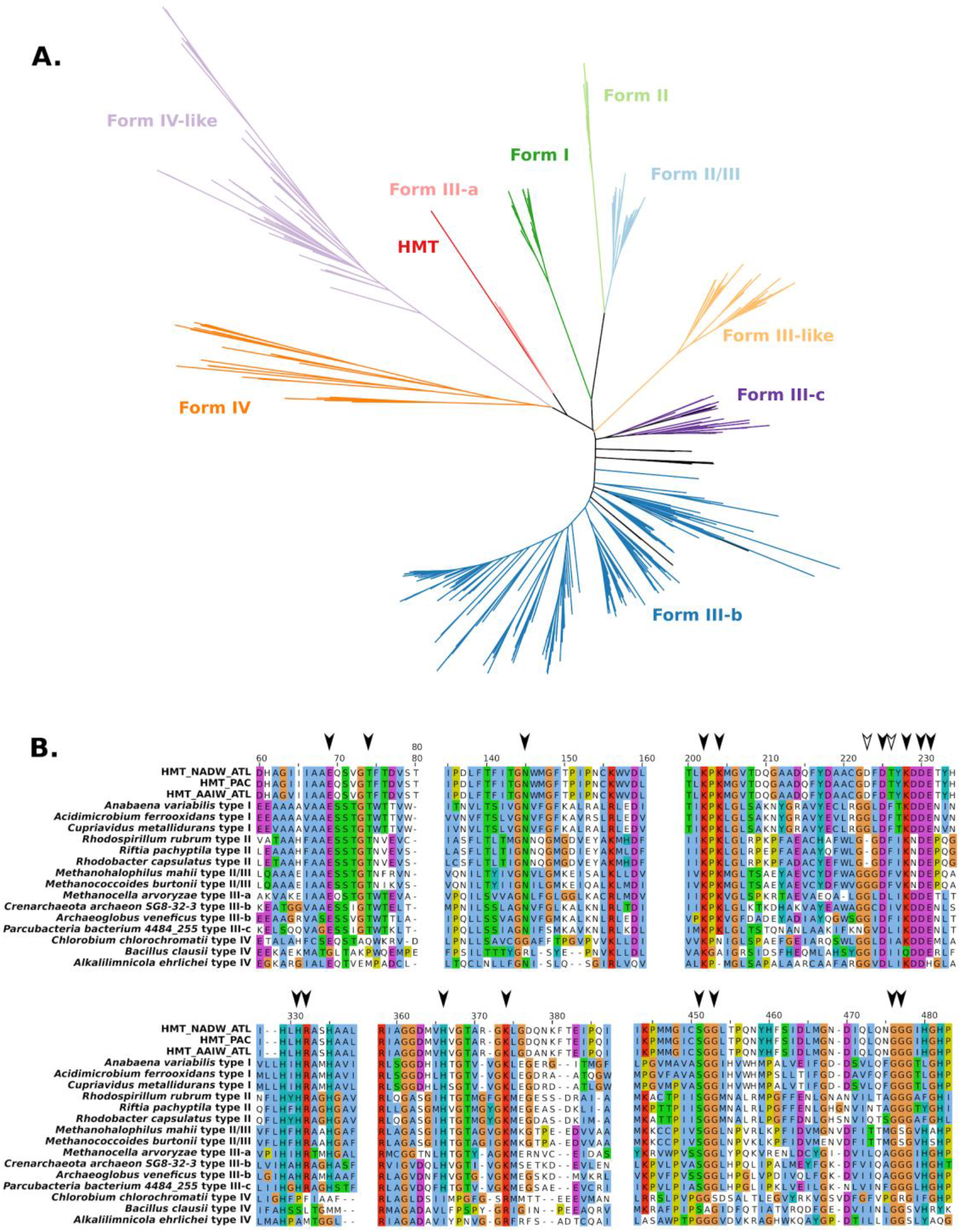
A) Phylogeny of RuBisCo large subunits. Reference sequences and RuBisCO form classifications were obtained from Jaffe & Banfield (2019). B) Sample alignment including different forms of RuBisCo, with arrows indicating 19 conserved residues shown to be important for enzymatic activity (see main text). The two unfilled arrows indicate residues that differ between HMT RuBisCo and other forms.

**Figure S5.**
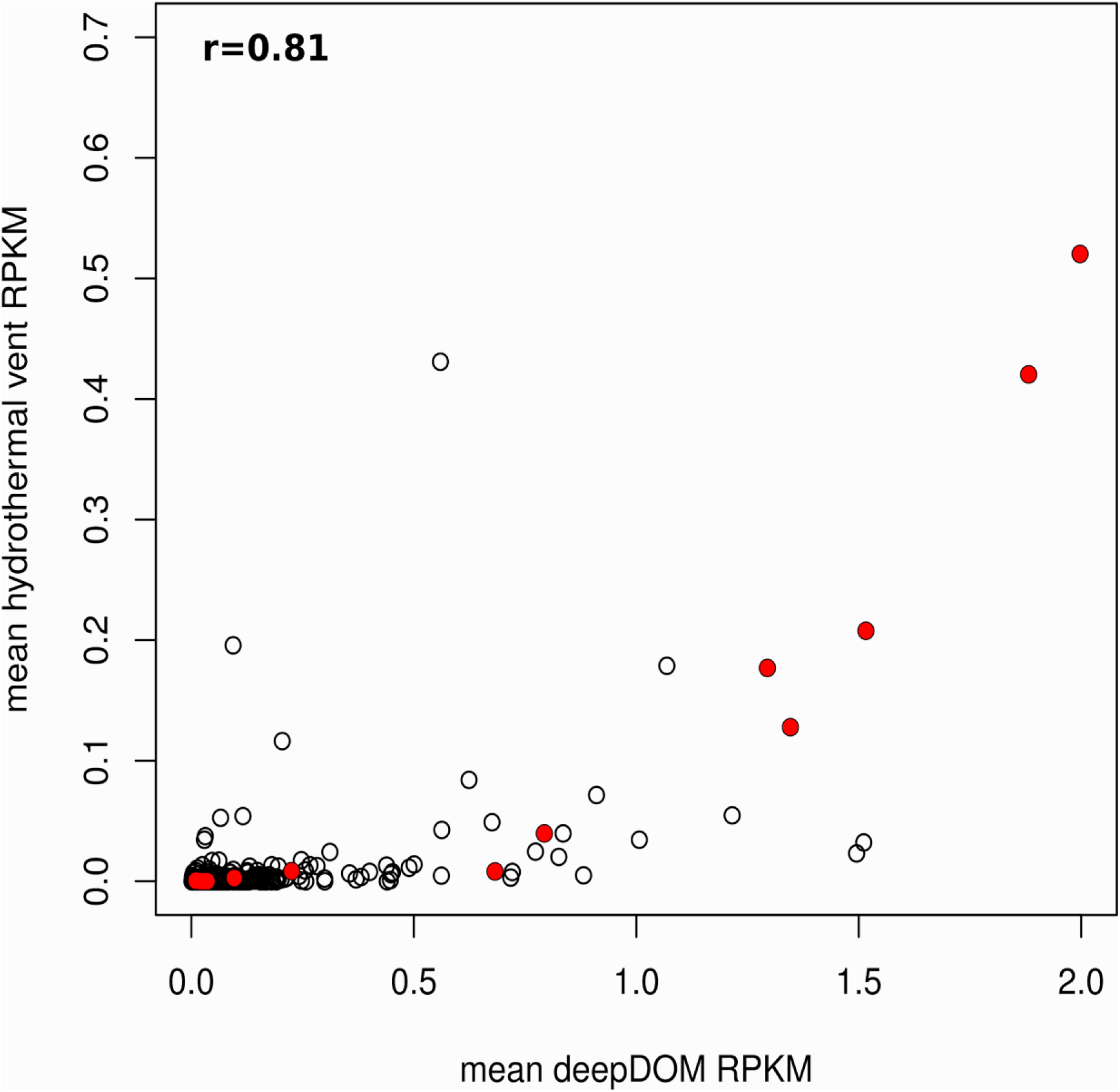
Comparison of the mean RPKM between the 10 DeepDOM metatranscriptomes and 12 metatranscriptomes associated with deep sea vents (55), indicating consistent transcriptional patterns of HMT across different marine habitats. PQQ-dehydrogenases are colored red. Both axes have been log10-transformed with pseudocounts added. The Pearson Correlation is provided on the top left.

## Supplementary Datasets

Dataset S1. Annotations for the orthologous groups generated in this study, as well as subsets for those identified as lost or gained in the HMT lineage.

Dataset S2. Annotations for the MAGs discussed in this study.

Dataset S3. Results for the metatranscriptome analyses performed in this study. Dataset S4. Read mapping statistics for the HMT relative abundance estimations.

